# Identification of a Female-produced Sex Attractant Pheromone of the Winter Firefly, *Photinus corrusca*

**DOI:** 10.1101/2022.12.07.519461

**Authors:** Sarah E. Lower, Gregory M. Pask, Kyle Arriola, Sean Halloran, Hannah Holmes, Daphné C. Halley, Yiyu Zheng, Douglas B. Collins, Jocelyn G. Millar

## Abstract

Firefly flashes are well-known visual signals used by these insects to find, identify, and choose mates. However, many firefly species have lost the ability to produce light as adults. These “unlighted” species generally lack developed adult light organs, are diurnal rather than nocturnal, and are believed to use volatile pheromones acting over a distance to locate mates. While cuticular hydrocarbons, which may function in mate recognition at close range, have been examined for a handful of the over 2000 extant firefly species, no volatile pheromone has ever been identified. In this study, using coupled gas chromatography - electroantennographic detection, we detected a single female-emitted compound that elicited antennal responses from wild-caught male winter fireflies, *Photinus corrusca*. The compound was identified as (1*S*)-*exo*-3-hydroxycamphor (hydroxycamphor). In field trials at two sites across the species’ eastern North American range, large numbers of male *P. corrusca* were attracted to synthesized hydroxycamphor, verifying its function as a volatile sex attractant pheromone. Males spent more time in contact with lures treated with synthesized hydroxycamphor than those treated with solvent only in laboratory two-choice assays. Further, using single sensillum recordings, we characterized a pheromone-sensitive odorant receptor neuron in a specific olfactory sensillum on male *P. corrusca* antennae and demonstrated its sensitivity to hydroxycamphor. Thus, this study has identified the first volatile pheromone and its corresponding sensory neuron for any firefly species, and provides a tool for monitoring *P. corrusca* populations for conservation, and further inquiry into the chemical and cellular bases for sexual communication among fireflies.

## Introduction

Summer firefly flashes are a nostalgic reminder of nature’s wonder. The bioluminescent signals of these charismatic beetles (family: Lampyridae), emitted from dusk into the night, are renowned for their function in mediating the finding, identifying, and choosing of mates (Lloyd 1966; Cratsley 2004). However, whereas all fireflies emit light in the larval stage, many firefly species have lost the ability to produce light as adults. These “unlighted” species generally lack developed adult light organs, are diurnally active, and are believed to use volatile pheromones acting over a distance to locate mates (McDermott 1964; Lloyd 1978, 1997; Ohba 1983; Branham and Wenzel 2001, 2003; De Cock and Matthysen 2005; Stanger-Hall et al. 2007; Stanger-Hall and Lloyd 2015; Martin et al. 2017).

Aside from the lack of light organs, evidence for diurnally active species’ use of volatile pheromones comes from (i) behavioral assays where males were attracted to a petri dish containing a female, sometimes even after the female had been removed (Lloyd 1972; De Cock and Matthysen 2005), (ii) comparative morphological studies where putative pheromone-using fireflies have smaller eyes and larger antennae than lighted species (Ohba 1978; Stanger-Hall et al. 2018), and (iii) comparison with a sister group - the click beetles (family: Elateridae), which diverged from fireflies ∼140-220 mya (Powell et al. 2022) and that generally use volatile pheromones to find mates (Tóth 2013; Serrano et al. 2018; Williams et al. 2019; Gries et al. 2021; Tolasch et al. 2022; Millar et al. 2022). While cuticular hydrocarbons (CHCs) that may function in mate recognition at close range have been identified for some species (Shibue et al. 2000, 2004; South et al. 2008; Ming and Lewis 2010), to date, no volatile pheromones of any firefly species have been characterized.

The winter firefly, *Photinus corrusca* Linnaeus (formerly *Ellychnia corrusca*; Zaragoza-Caballero et al. 2020), is an unlighted firefly native to North America, ranging from Alaska to Mexico (Fallon et al. 2021, Figure 1). Adult *P. corrusca* completely lack light organs and are diurnal. In contrast to familiar summer-active species, *P. corrusca* emerge as reproductively immature adults in the fall, overwinter on the sides of large-diameter trees, and mature their ovaries and testes in the spring for mating in the late spring/early summer (Rooney and Lewis 2000; Faust 2012). Based on behavioral observations and comparison with related firefly species, males are likely the searching sex, flying toward perched calling females, landing nearby, and scrambling to approach (Lloyd 1972; Vencl and Carlson 1998; De Cock and Matthysen 2005). Chemical signals are likely important for *P. corrusca* mating behaviors because the male, upon contact, will vigorously antennate and pass his palps over the female while initiating a two-stage copulation that may last over 20 h (Rooney and Lewis 2000).

**Fig. 1.**
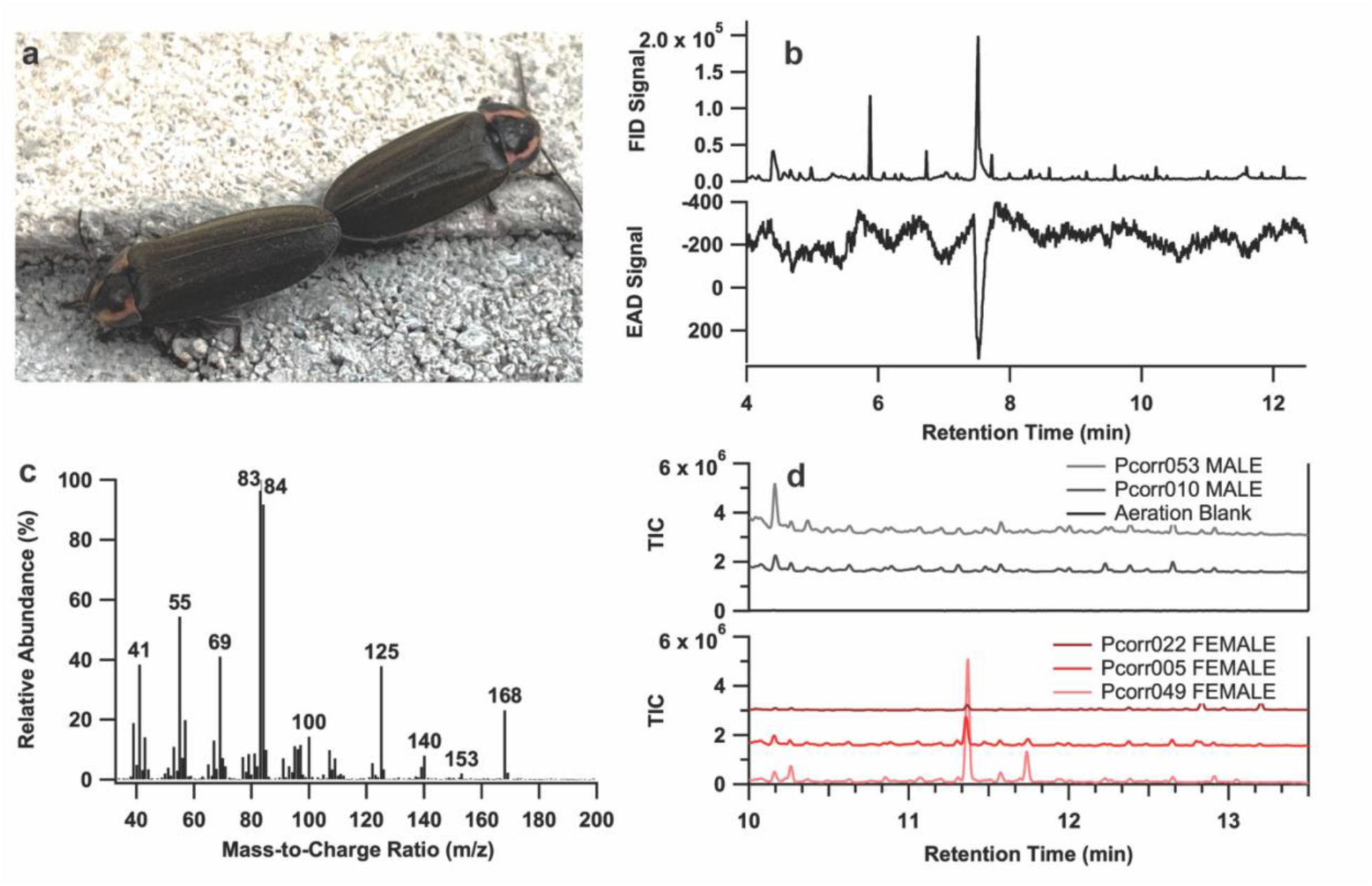
A female-emitted *Photinus corrusca* compound elicited an antennal response from a male *P. corrusca*. (**a**) A pair of *P. corrusca* individuals engaged in mating in Middlebury, VT. Photo: G. Pask. (**b**) Coupled GC-FID and GC-EAD chromatograms showing the response of an antenna of a *P. corrusca* male to the extract of the headspace odors collected from live *P. corrusca* females. Top trace = GC-FID response, inverted trace = antennal response. (**c**) EI mass spectrum of the unknown compound produced by female *P. corrusca* [M]^+^ = 168. (**d**) GC-MS total ion chromatograms (TIC) for headspace samples from male (top) and female (bottom) *P. corrusca* individuals collected in Pennsylvania in 2022. The compound of interest is observed at RT = 11.36 min. The structurally-related camphorquinone was at RT = 11.74 min. A headspace collection blank is shown and did not indicate a signal at retention times of interest. Pcorr### SEX - unique identifier of an assessed individual

Further, laboratory behavioral assays have demonstrated that males do not attempt to copulate with freeze-killed females that have had their CHCs and other surface chemicals extracted with solvent (South et al. 2008; Ming and Lewis 2010). However, the precise compound(s) involved in these close-range interactions remain unknown. Ming and Lewis (2010) identified a female-specific compound, sampled from *P. corrusca* elytra using either solid-phase microextraction (SPME) or a hexane rinse, as *exo,exo*-2,3-camphanediol. However, the synthetic racemate of this compound did not elicit attraction or mating behavior in laboratory behavioral assays. Additionally, these previous studies focused on close-range, contact interactions, such as those mediated by CHCs, which are relatively nonvolatile, making them unlikely candidates for long-range attractant pheromones (Blomquist et al. 2020).

Here, using coupled gas chromatography - electroantennographic detection (GC-EAD) and coupled gas chromatography-mass spectrometry (GC-MS), we identified a female-produced compound from headspace volatiles of female *P. corrusca* that elicited responses from antennae of male beetles. The compound was synthesized and tested in field trials at two sites across *P. corrusca*’s range, in Vermont and Pennsylvania, and in laboratory assays in Pennsylvania. Finally, using scanning electron microscopy (SEM) and single sensillum recordings (SSR) we located an odorant receptor neuron (ORN) in males that responds specifically to this compound. To our knowledge, this is the first sex attractant pheromone identified from any firefly species.

## Materials and Methods

### Specimen Collection and Maintenance

Adult male and female *P. corrusca* were collected during March and April 2021 on the sides of large-diameter trees in Lewisburg, PA (Lat: 40°55’ N, Lon: 76°54’ W). At this point in the season they were likely emerging from winter diapause (Rooney and Lewis 2000). Because *P. corrusca* do not mate until later in the season, after they have matured their ovaries and testes (Rooney and Lewis 2000), individuals were presumed to be unmated upon collection. Adult *P. corrusca* were easily identified, as they were the only beetle found on the sunny side of large diameter trees in that area at that time of year. To prevent mating, after capture individuals were housed individually in disposable polypropylene narrow *Drosophila* vials (Genesee Scientific; Flystuff #32-120; El Cajon CA, USA) sealed with a reusable foam plug (Genesee Scientific; Flystuff #59-200; El Cajon CA, USA). Because winter fireflies are one of the few firefly species that are suspected to feed as adults, potentially consuming nectar or sap (Rooney and Lewis 2000), each vial was furnished with a ⅓ piece of cotton dental lozenge (Richmond Dental & Medical #200204; Seattle WA, USA) soaked in 10% honey water that was replaced weekly to prevent microbial growth. To maintain humidity and simulate natural conditions, vials were maintained in an Adaptis Conviron A1000 Growth Chamber (Conviron US; Pembina ND, USA) at 80-90% humidity, which was adjusted monthly to match the average day length and average high temperature for each month for Lewisburg, PA (WeatherSpark; Supplementary Table S1). Where average high and/or low temperatures were outside the limits of chamber specifications, the minimum temperature was set to 12°C. Individuals were periodically removed from their vials and assessed for mating receptiveness by observing female and male behavior when they were placed together in a BugDorm® cage (BioQuip; BugDorm-1 Insect Rearing Cage; Rancho Dominguez CA, USA). When successful attraction and mating in the BugDorm® cage were observed in May, remaining putative virgin males and females were shipped in their individual vials in double containment overnight to the University of California Riverside quarantine facility (USDA-APHIS-PPQ permit #P526P-20-02507) for collection of headspace volatiles and analyses of the resulting extracts.

### Acquisition of Firefly-Produced Compounds

To capture firefly-emitted compounds, collections of headspace volatiles were conducted using wide-mouth 250 mL Teflon jars (Thermo Fisher Scientific; #24030250; Hampton NH, USA), with the screw cap lids fitted with Swagelok bulkhead unions (Swagelok; Solon OH, USA) to connect inlet and outlet tubes. Air, purified by passage through granulated activated charcoal (14-16 mesh; Fisher Scientific), was pulled through the system by vacuum at 250 mL/min. Volatiles were adsorbed onto ∼50 mg of thermally desorbed activated charcoal (50-200 mesh; Fisher Scientific) held between glass wool plugs in a short piece of glass tubing. Aerations were conducted under ReptiSun 100 UVB lights (Zoo Med Laboratories Inc.; San Luis Obispo CA, USA) at 24°C with a 16:8 h L:D cycle to simulate summer daytime conditions. Volatiles were eluted from the charcoal with 500 µL of dichloromethane (CH_2_Cl_2_).

All fireflies were aerated individually and given two vials with wicks containing either water or 10% honey solution. Earlier aerations of fireflies were run over 3-5 d and then increased up to 8 d for later aerations. Individuals were aerated repeatedly until death, from May 13 – June 14, 2021. A total of seven females and four males were aerated, yielding 24 extracts of females and 5 of males (Supplementary Table S2). To further examine sex-specific differences in headspace-sampled compounds, an additional group of male and female fireflies were collected February - May, 2022 (Supplementary Table S3) and aerated individually in Lewisburg, PA between May 8 - June 21, 2022 following identical protocols, except the air temperature was held at approximately 22°C and light was provided by a northwest-facing window. A subset of assessed individuals with relatively intact carcasses are retained in 95 - 100% EtOH at -80°C in the SEL molecular collection at Bucknell University (Supplementary Tables S2 - 3).

### Chemical Analysis

To identify firefly-emitted volatile compounds, the aeration extracts were initially analyzed by GC-MS in splitless mode with an Agilent 7820A GC interfaced to an Agilent 5977E mass selective detector (Agilent Technologies; Santa Clara CA, USA). A DB-5MS column was used (30m +10m Duraguard x 0.25 mm ID x 0.25 micron film; J&W Scientific; Folsom CA, USA), with a temperature program of 40°C for 1 min, then ramping at 10°C /min to 280°C, followed by a hold for 10 min. The injector and transfer line temperatures were 250°C and 280°C, respectively. Mass spectra were collected in electron ionization mode (EI) at 70 eV, with a scan range of *m/z* 30-500. Samples for sex-specific volatile compound emission follow-up were analyzed at Bucknell University using the same method but a slightly different hardware configuration (8860/5977B GC-MS, Agilent Technologies) and column (HP-5ms-UI; 30 m x 0.25 mm ID x 0.25 micron film; J&W Scientific).

Subsequently, aeration extracts and synthetic standards were analyzed by GC-EAD with an HP 5890 Series II GC equipped with a DB-5MS Column (30m + 10m retention gap x 0.25mm ID x 0.25 micron film; Agilent Technologies). The injector and detector temperatures were 250 °C and 280°C, respectively, with a column head pressure of 200 kPa. The temperature program was 50°C for 1 min, then ramping at 20°C/min to 280°C, then held for 10 min. The column effluent was split to the flame ionization detector (FID; 280°C) and a heated transfer line (260°C) with a glass X-cross, with additional helium being added at the fourth arm at 3 ml/min as makeup gas in order to maintain the flow rate to both the FID and EAD. Effluent passing through the transfer line was directed into a humidified airstream at 650 ml/min in a glass tube (15 mm ID), which then passed over the *P. corrusca* antennal preparation.

Antennae of *P. corrusca* were prepared by removing an entire antenna with a razor blade followed by removing a portion of the distal tip. The antenna was then placed between two glass capillary electrodes filled with saline solution (7.5 g NaCl, 0.21 g CaCl_2_, 0.35 g KCl, and 0.20 g NaHCO_3_ in 1 L Milli-Q purified water). Electrical connection was made with a 0.2 mm diameter gold wire in each capillary, attached to a custom-built amplifier. Signals from the FID and the amplifier were recorded in tandem using Peak-Simple software (SRI International; Menlo Park CA, USA). All GC-MS and GC-EAD files derived from sample aerations are available on Figshare (DOI:10.6084/m9.figshare.21648497).

To isolate the compound of interest, five aeration extracts from females in CH_2_Cl_2_ (∼2.5 mL) were first combined and concentrated to ∼0.1 mL under a gentle stream of nitrogen, and 0.5 mL pentane was added. The sample was blown down again, and the procedure was repeated to remove as much of the CH_2_Cl_2_ as possible. The concentrated sample in ∼0.1 mL pentane was then loaded onto a column of silica gel (230-400 mesh, 100 mg), pre-wetted with pentane. The column was eluted sequentially with 2 × 0.5 mL pentane, 2 × 0.5 mL 25% ether in pentane, and 2 × 1 mL ether, collecting each as a separate fraction. The compound of interest with an apparent molecular ion at *m/z* 168 started eluting in the 2nd 25% ether fraction (Frac. 4), with the bulk eluting in the first 100% ether fraction.

To aid in the identification of the compound, a series of microderivatization experiments were conducted. An aliquot of the 100% ether fraction was diluted 1:1 with pentane, ∼ 1 mg of 10% Pd on carbon was added, and the mixture was stirred under H_2_ for 1 h. The mixture was then filtered through a small pad of celite, rinsing with ether, concentrated to ∼100 µL, and an aliquot was analyzed by GC-MS. Approximately half of the remaining H_2_-reduced sample was diluted with 0.5 mL ether, and ∼ 2 mg LiAlH_4_ was added. The mixture was stirred for 1 h at room temp, then quenched by careful addition of 0.1 mL aqueous 1 M HCl, followed by 0.5 mL saturated brine. The mixture was vortexed, and the ether layer was removed and dried over anhydrous Na_2_SO_4_, and an aliquot was analyzed by GC-MS. To test the effects of oxidation, a second aliquot of fraction 4 (eluted with 25% ether in pentane) was concentrated to ∼ 10 μL, then diluted with 0.1 mL CH_2_Cl_2_, ∼2 mg finely powdered pyridinium dichromate was added, and the mixture was stirred for 1 h at room temp. The mixture was then diluted with 0.5 mL ether, and filtered through a plug of celite. An aliquot was analyzed by GC-MS.

### Syntheses of Pheromone Candidates

All solvents were Optima grade (Fisher Scientific) unless otherwise noted. Anhydrous diethyl ether stabilized with BHT was purchased from Fisher Scientific. Solutions of crude reaction products were dried over anhydrous Na_2_SO_4_ and concentrated by rotary evaporation under partial vacuum. Flash and vacuum flash chromatography were carried out on silica gel (230– 400 mesh; Fisher Scientific). TLC analyses were conducted on aluminum-backed sheets of analytical silica gel 60 F254 (Merck, Darmstadt, Germany), and compounds were visualized by spraying with 10% phosphomolybdic acid in ethanol and heating. Yields are reported as isolated yields of chromatographically pure products unless otherwise noted. Mass spectra were obtained with an HP 6890 GC (Agilent Technologies) equipped with a DB-17 column (30 m × 0.25 mm × 0.25 micron film; J&W Scientific), coupled to an HP 5973 mass selective detector in EI mode (70 eV) with helium carrier gas. Purity was assessed by gas chromatography with an HP 5890 GC equipped with a DB-5 column (30 m × 0.25 mm × 0.25 micron film), unless otherwise noted. Isomeric and enantiomeric purity were assessed by gas chromatography with an HP 5890 GC equipped with a chiral stationary phase β-cyclodextrin column (Cyclodex-B, 30 m × 0.25 mm × 0.25 µm film + 2 m × 0.25 mm ID deactivated fused silica; J&W Scientific). Nuclear magnetic resonance (NMR) spectra were recorded as CDCl_3_ solutions on either a Bruker Avance 500 or Bruker NEO 400 spectrometer. Chemical shifts are reported in ppm relative to CDCl_3_ (^1^H 7.26 ppm; ^13^C 77.0 ppm).

### Partial Reduction of (1*S*)-Camphorquinone with LiAlH_4_

LiAlH_4_ (7.5 mg, 0.2 mmol) was added to a solution of (*S*)-(-)-camphorquinone (166 mg, 1 mmol, TCI Americas, Portland OR, USA) in 2.5 mL dry ether. The mixture was stirred 1 h at room temp, then quenched with 1 mL of 1M HCl. The mixture was then diluted with 10 mL brine and extracted with ether. The ether solution was dried and concentrated, and the residue was fractionated by flash chromatography on silica gel, eluting with 25% EtOAc in hexane, achieving a partial separation of the isomers. The fractions were checked by GC on a DB-5 column, matching the retention time of one hydroxycamphor isomer with that of the insect-produced compound. An aliquot was then reanalyzed by GC-MS on a DB-17 column, matching both the retention time and the mass spectrum of one of the isomers with those of the insect-produced compound.

### Reduction of (1*S*)-Camphorquinone with NaBH_4_

The following procedure was adapted from Xu et al. (2006). Identities of each product isomer were verified by published ^1^H-NMR spectral data (Xu et al. 2002; Neisius and Plietker 2008).

A dry flask flushed with Ar was charged with (1*S*)-camphorquinone (83.1 mg, 0.50 mmol, 3.8 eq; TCI Americas, Portland OR, USA), diethyl ether (0.5 mL), and methanol (0.5 mL). The flask was cooled to 0ºC followed by the addition of NaBH_4_ (5 mg, 0.132 mmol, 1 eq) in one portion. The reaction was stirred at 0ºC for 30 min, then quenched with water (167 µL) and diluted with brine (300 µL). The mixture was extracted with diethyl ether (3×1 mL), and the organic phases were combined, washed with brine, dried over anhydrous Na_2_SO_4_, and concentrated under reduced pressure to yield a crude white solid composed of a 65:35 mixture of isomers. This crude material was purified by vacuum flash column chromatography (20% EtOAc/Hex) to yield 59 mg (73%) of a 65:35 mixture of isomers (1*S*)-*exo*-**2** and (1*S*)-*exo*-**3**. (1*S*)-*exo*-**2**: ^1^H NMR (500 MHz, CDCl_3_): δ 3.73 (s, 1H), 3.23 (bs, 1H), 2.07 (d, *J* = 4.9 Hz, 1H), 1.97 (dddd, *J* = 12.4, 4.8 Hz, 1H), 1.63 (ddd, *J* = 12.4, 3.7, 1H), 1.47-1.30 (m, 2H), 0.96 (s, 3H), 0.92 (s, 3H), 0.90 (s, 3H). ^13^C NMR (126 MHz, CDCl_3_) δ 220.52, 77.26, 57.02, 49.22, 46.72, 28.52, 25.11, 20.95, 20.00, 8.96. (1*S*)-*exo*-**3**: ^1^H NMR (500 MHz, CDCl_3_): δ 3.53 (s, 1H), 3.23 (bs, 1H), 2.14 (d, 1H), 1.89 (m, 1H), 1.82 (m, 1H), 1.47-1.30 (m, 2H), 1.01 (s, 3H), 1.00 (s, 3H), 0.90 (s, 3H). ^13^C NMR (126 MHz, CDCl_3_) δ 219.27, 79.35, 58.58, 49.15, 46.51, 33.77, 21.12, 20.28, 18.78, 10.25. (1*S*)-*exo*-**2** and (1*S*)-*exo*-**3**: EI-GC/MS *m/z* (%): 168 (M^+^, 36), 153 (6), 140 (9), 125 (41), 107 (11), 100 (13), 84 (92), 83 (100), 71 (22), 70 (33), 69 (42), 57 (20), 55 (53), 43 (22), 41 (46).

### Reduction of (1*S*)-Camphorquinone with Zn and AcOH

The following procedure was adapted from Hückel and Fechtig (1962) with additional insights from Templeton et al. (1990). Identities of each product isomer were verified by comparison with published ^1^H-NMR spectral data (Tan et al. 2011). A flask was charged with acetic acid (200 µL, 3.50 mmol, 5.8 eq), (1*S*)-camphorquinone (100 mg, 0.602 mmol, 1 eq), and water (1.8 mL) and heated to 100 ºC. The heterogenous yellow mixture was briefly stirred, then zinc dust (100 mg, 1.53 mmol, 1.5 eq) was added in portions over 1 min. The reaction was stirred for 20 min at 115 ºC, becoming colorless. The reaction was allowed to cool, then diluted with water (2 mL) and extracted with diethyl ether (2.5 mL). The aqueous layer was extracted with additional diethyl ether (2.5 mL), and the combined organic layer was sequentially washed with sat. aq. NaHCO_3_ (1 mL) and brine (1 mL), dried, concentrated, and the crude material was purified by vacuum flash column chromatography (20% EtOAc/Hex) to yield 81 mg (80%) of a 56:44 mixture of the two *endo* isomers (1*S*)-*endo*-**2** and (1*S*)-*endo*-**3**. (1*S*)-*endo*-**2**: ^1^H NMR (500 MHz, CDCl_3_): δ 4.20 (d, 1H), 3.35 (bs, 1H), 2.26 (t, *J* = 4.0 Hz, 1H), 1.93 (m, 1H), 1.70 (m, 1H), 1.67 (m, 1H), 1.38 (m, 1H), 0.98 (s, 3H), 0.90 (s, 3H), 0.85 (s, 3H). (1*S*)-*endo*-**3**: ^1^H NMR (500 MHz, CDCl_3_): δ 3.84 (s, 1H), 3.35 (bs, 1H), 2.24 (dd, *J* = 5.4, 1H), 1.96 (m, 2H), 1.38 (m, 2H), 1.03 (s, 3H), 0.95 (s, 3H), 0.90 (s, 3H). (1*S*)-*endo*-**2** and (1*S*)-*endo*-**3**: ^13^C NMR (126 MHz, CDCl_3_) δ 220.95, 219.53, 78.73, 74.40, 59.40, 58.41, 50.22, 48.58, 43.08, 43.01, 32.48, 24.99, 24.62, 19.94, 19.32, 18.76, 17.84, 12.91, 9.26. EI-GC/MS *m/z* (%): 168 (M^+^, 68), 153 (14), 139 (11), 135 (7), 125 (32), 109 (14), 107 (11), 95 (29), 84 (75), 83 (89), 71 (63), 70 (100), 69 (52), 55 (50), 44 (40), 43 (67).

### Reduction of (1*S*)-Camphorquinone with L-Selectride

The following procedure was adapted from Kouklovsky et al. (1990). A dry flask flushed with Ar was charged with (1*S*)-camphorquinone (5.83 g, 35.1 mmol, 1 eq) and dry THF (200 mL), then cooled to -78 ºC. L-Selectride (1 M in THF, 42 mmol, 1.20 eq; MilliporeSigma, St. Louis MO, USA) was added over 1 h, and the mixture was stirred an additional 5 min. The reaction was quenched by addition of 3 M HCl in methanol (16 mL) over 10 min at -78 ºC, then allowed to warm to room temperature. The resulting mixture was diluted with brine (500 mL) and the products were extracted with dichloromethane (4 × 200 mL). The combined organic layers were dried, concentrated, and purified by vacuum flash chromatography (5% - 25% EtOAc/Hexanes). The fractions rich in the desired product were concentrated and subjected to flash column chromatography (20% EtOAc/Hexanes) to partially separate the isomers of hydroxycamphor.

The resulting material was sublimed with a Kugelrohr distillation apparatus (55 ºC, 0.1 torr), yielding 397 mg (7%) of (1*S*)-*exo*-**2**, 92% isomerically pure (NMR). Impure fractions were consolidated and purified again by flash chromatography to yield an additional 298 mg of (1*S*)- *exo*-**2**, ∼95% isomerically pure. ^1^H NMR (500 MHz, CDCl_3_) δ 3.72 (s, 1H), 3.28 (bs, 1H), 2.06 (d, *J* = 4.7 Hz, 1H), 2.02 – 1.91 (dddd, 1H), 1.62 (ddd, 1H), 1.46 – 1.29 (m, 2H), 0.96 (s, 3H), 0.91 (s, 3H), 0.89 (s, 3H). ^13^C NMR (126 MHz, CDCl_3_) δ 220.51, 77.26, 57.01, 49.22, 46.71, 28.51, 25.10, 20.94, 19.98, 8.95. EI-GC/MS *m/z* (%): 168 (M^+^, 32), 153 (2), 140 (9), 125 (42), 107 (10), 100 (14), 84 (92), 83 (100), 69 (38), 57 (19), 55 (51), 43 (14), 41 (37).

### Davis Oxidation of Camphor

The following procedure was adapted from Davis et al. (1984). This paper reported the reaction product as *endo*-hydroxycamphor *endo*-**2**. However, in our hands, and based on other published reports (Neisius and Plietker 2008; Piątek and Chapuis 2021) this reaction yielded *exo*-hydroxycamphor *exo*-**2** as the major product. Our results were corroborated with NMR spectral data.

In a preliminary synthesis, a 2:1 mixture of (1*R*)- and (1*S*)-camphor (0.33 mmol) was used in the Davis oxidation with the intention of producing a differentiable mixture of *endo*-hydroxycamphor enantiomers. After discovering that the insect-produced compound and the minor enantiomer from the Davis oxidation product matched, a larger scale reaction from (1*S*)-camphor (>98%) was run as follows. A dry flask flushed with Ar was charged with THF (40 mL) and potassium hexamethyldisilazide (KHMDS 1 M in THF, 2.22 mL, 2.22 mmol, 1 eq), then cooled to -78 ºC. (1*S*)-Camphor (338 mg, 2.22 mmol, 1 eq; TCI Americas) was added in portions over 1 min and the mixture was stirred 30 min at -78 ºC. A solution of the Davis reagent, 2-(phenylsulfonyl)-3-phenyloxazaridine (881 mg in 15 mL THF, 3.37 mmol, 1.5 eq; Enamine, Monmouth Junction NJ, USA) was added dropwise over 20 min. The reaction was stirred at -78 ºC for 55 min. Upon completion, the reaction was quenched with sat. aq. NH_4_Cl (10 mL) at -78 ºC and allowed to warm to room temperature. Brine (30 mL) and diethyl ether (30 mL) were added and the layers were separated. The aqueous layer was reextracted with diethyl ether (30 mL), then the combined organic layers were washed with brine (20 mL), dried, and concentrated. The resulting cloudy yellow oil was dissolved in 4:1 dichloromethane:hexanes and partially purified by vacuum flash column chromatography (20% EtOAc/hexanes). The fractions enriched in the desired product were concentrated and repurified by flash column chromatography (25% EtOAc/hexanes). The product was then purified further by Kugelrohr distillation (80 ºC, 1.85 torr) to yield 182 mg of product (49%), containing ∼85% of the desired insect-produced exo-hydroxycamphor isomer (1*S*)-*exo*-**2** and 15% of the *endo*-hydroxycamphor isomer (1*S*)-*endo*-**2**. EI-GC/MS *m/z* (%): 169 (4), 168 (M^+^, 34), 153 (2), 140 (9), 125 (43), 107 (11), 100 (15), 84 (93), 83 (100), 69 (38), 57 (18), 55 (50), 43 (14), 41 (35).

### Field Trials

Field trials were conducted at two sites: Sundance Ridge in Lewisburg, PA (May 13 -14 2022, 40.91897° N, 76.90109° W) and Middlebury College in Middlebury, VT (May 20 -21 2022, 44.01365° N, 73.18275° W). At each site, 10 sticky traps were deployed every 20 m in a single transect, with the sticky surface south-facing. Sticky traps consisted of packing-tape covered 10-inch cardboard cake rounds secured to the top of a 4-foot fence post (Supplementary Information Figure S1). Vertical sticky traps, instead of horizontal, were used to mimic natural *P. corrusca* mating conditions on the sides of trees. Lures were made by dosing 8 mm red rubber septa (Ace Glass; Vineland NJ, USA) with 100 uL of a 10 mg/mL solution of 92% pure synthetic pheromone in pentane (1 mg pheromone). Controls consisted of septa dosed with 100 uL pentane. Control and experimental lures were held in separate glass bottles at -20℃ until field deployment. Lures were deployed inside a wire mesh tea strainer suspended from a paper-clip hook in the middle of the cake round. A coat of TangleTrap (Tanglefoot, Amazon) was applied to the remaining cake round surface (front side only) to provide a sticky surface in which attracted males, landing nearby and scrambling toward the pheromone source, would adhere. To avoid potential bias in TangleTrap application based on trap treatment status, the sticky coating was applied prior to trap assignment as either a control (solvent only) or experimental (solvent + pheromone) trap. Treatment (control versus experimental) of the first trap in the transect was determined using a random number generator (https://www.random.org/; even = experimental), and the status alternated along the transect. After 24 h, the individuals stuck in each trap were removed, preserved in 95% EtOH, and identified to sex by examination of external morphology under a dissecting microscope (Leica EZ4; Supplementary Information Figure S2). The PA population samples are stored in the -80℃ Lower molecular collection at Bucknell.

### Laboratory Assays

Two-choice assays were conducted between control (solvent-only) and experimental (solvent + pheromone) septa in laboratory assays with male *P. corrusca* collected in Lewisburg, PA from February 2, 2022 to May 5, 2022 (Supplementary Table S4) and maintained as described earlier. Assays were conducted in a 9 cm glass petri dish rinsed and air-dried three times with isopropanol to remove contaminants. Once dry, the dish was laid over a paper template marking rubber septa placements, labeled A or B. Septa were prepared as in field trials by applying 100 uL of either pentane (control) or 100 uL of 10 mg/mL hydroxycamphor dissolved in pentane (experimental) to the cup of the septum and allowing the solvent to evaporate. Septum placement for each trial was randomized using a random number generator (even = experimental as A; odd = control as A). Assays were recorded with a PiSpy, an automated video recording device that utilizes a Raspberry Pi computer along with LEDs for red and white light cycling (Morris et al. 2022). The PiSpy recording device was set to record for 16 min with the arena illuminated by two white LED printed circuit boards and diffusers (Morris et al. 2022). A single, male *P. corrusca* was then placed in the center of the arena and a box lined with aluminum foil was placed over the setup for better light diffusion for image quality. To accommodate an adjustment period, only the last 15 min after specimen placement were analyzed. During the observation period, the duration of time spent either touching (any part of body in contact with septa), mounting (all legs on septa), or copulating (attempted aedeagus insertion), as well as total time in contact (sum of touching, mating, and copulation), was recorded for both the control and experimental septa.

### Statistical Analysis

For field bioassays, because catch data were not normally distributed, the attractive effect of the pheromone relative to solvent controls was assessed using a one-sided Mann Whitney Wilcoxon test in R (R Core Team 2020). Locations were tested separately because there may be differences in phenology between the field sites due to latitude (Faust 2012). For laboratory assays, the difference in total time in contact with either control or experimental septa was assessed with paired Wilcoxon rank-sum tests. All R scripts and outputs are hosted on GitHub (https://selower.github.io/Winter_firefly_pheromone_project/).

### Scanning Electron Microscopy (SEM)

From *P. corrusca* collected from Middlebury, VT in May-June, antennae were removed while under CO_2_ anesthesia and then dried sequentially with hexanes, acetone, and 95% ethanol in a watch glass. Dried antennae were mounted on an aluminum specimen mount and coated with gold-palladium using an EffaCoater Au-Pd Sputter Coater (Ernest Fullam Inc.; Latham NY, USA). Coated antennae were then imaged using a Vega 3 LMU Scanning Electron Microscope (Tescan; Brno, Czech Republic) with an accelerating voltage of 5 mV and a beam intensity of 8.

### Single Sensillum Recording (SSR)

Adult male *P. corrusca* were collected in pheromone-baited traps in Middlebury, VT in May 2022. Collected fireflies were housed under ambient light conditions in plastic vials with a moistened Kimwipe 1-4 d before being prepared for electrophysiology. Fireflies were immobilized on a microscope slide with the ventral side up using double-sided tape. Additional strips of double-sided tape were used to immobilize the base and distal ends of the antennae, leaving both the head and middle segments accessible to electrodes. The preparation was mounted under a BX51WI microscope (Olympus; Center Valley PA, USA) and kept under a continuous 20 mL/s flow of humidified air using a CS-55 stimulus controller (Syntech; BuchenBach, Germany). Sharp glass electrodes filled with sensillum lymph Ringers solution (Kaissling 1995) and a Ag-Cl wire were inserted sequentially into the head (reference electrode) and the sensillum of interest (recording electrode) using micromanipulators. Specific sensilla were initially identified by morphology, spontaneous firing rate, and number of neurons and subsequently analyzed from the recorded data. Extracellular recordings were amplified using a Model 3000 AC/DC Differential Amplifier and Headstage (A-M Systems; Carlsborg WA, USA) and digitized using a Digidata 1550B Data Acquisition System (Molecular Devices; San Jose CA, USA). Data were sampled at 10 kHz and AC filtered at 10-1,000 Hz. No more than two sensillum recordings were obtained per firefly to limit potential desensitization of the olfactory neurons. Carcasses are retained in the -80℃ archival Lower collection at Bucknell University.

To test the detection limits of pheromone-sensitive sensilla, different doses of pheromone were delivered to the sensillum using cartridges made from glass Pasteur pipettes and 1 mL pipette tips. Pheromone solutions in pentane (pheromone doses of 0.1 - 100 µg) or pentane control were applied to the inside of a Pasteur pipette, which was left open for 5 min to allow for pentane evaporation before being sealed with a 1 mL pipette tip and parafilm squares at either end. Pheromone cartridges were inserted into the continuous stream of humidified air and 16 mL/s charcoal-filtered air switched from the blank cartridge to the pheromone cartridge for 1 s using the CS-55 stimulus controller. Each recording consisted of multiple 10 s traces (1 s of baseline firing, 1 s of firing during pheromone stimulus, and 8 s of post stimulus firing).

Neuronal responses to pheromone stimulation were collected and analyzed manually using Axoscope 10 (Molecular Devices). The baseline firing rate was determined for each neuron during the 1 s before each stimulus, and the stimulus firing rate was counted in a 500 ms window between 150-650 ms of the stimulus delivery to account for the delay in the airflow delivery system. The change in firing frequency (∆ spikes/s) was calculated by subtracting the baseline firing frequency from that of the stimulus. Data from 8 basiconic sensilla from 5 *P. corrusca* individuals were analyzed and plotted using Prism 9 (Graphpad; San Diego CA, USA).

## Results

### Identification of the Pheromone Candidate

A sex-specific compound was reproducibly observed in extracts of headspace odors sampled from live female *P. corrusca* fireflies collected from February through June of 2021 and 2022 in Union and Montour Counties, Pennsylvania (Figure 1). This compound, and no others in the extracts, elicited consistent, strong responses from the antennae of male fireflies by coupled gas chromatography-electroantennogram detection (GC-EAD) (Figure 1), suggesting that it was a likely candidate for a sex pheromone component. When analyzed by coupled gas chromatography-mass spectrometry with electron ionization (GC-EI-MS), the compound had an apparent molecular ion ([M^+^]) at *m/z* 168 (Figure 1c), for possible molecular formulae of C_12_H_24_, C_11_H_20_O, or C_10_H_16_O_2_, with 1, 2, or 3 sites of unsaturation, respectively.

When a crude extract was fractionated by liquid chromatography on silica gel, the compound eluted with 25-100% ether, indicating that it was of medium polarity. This eliminated the possibility of the molecular formula being C_12_H_24_, i.e., a hydrocarbon with no polar functional groups. Several microderivatization experiments were useful in providing additional information: (i) the unknown was unchanged by catalytic hydrogenation, indicating the absence of carbon-carbon pi bonds; (ii) the unknown seemed to be reduced with LiAlH_4_, such that the unknown peak disappeared from the GC trace, but the product(s) were not found; and (iii) when an aliquot of the insect extract was oxidized with pyridinium dichromate (PDC, Figure 2a), the unknown disappeared and the peak due to a minor compound in the crude extract that had been tentatively identified as camphorquinone **1** (*m/z* 166) by a match with the NIST mass spectral database, increased in size. The identification was confirmed by matching the retention time and mass spectrum with those of an authentic standard of camphorquinone **1**. The facts that the unknown had a molecular weight that was two mass units higher than that of camphorquinone **1**, and that PDC oxidation converted the unknown to camphorquinone **1**, indicated that the unknown had to be a hydroxycamphor isomer (Figure 2).

**Fig. 2.**
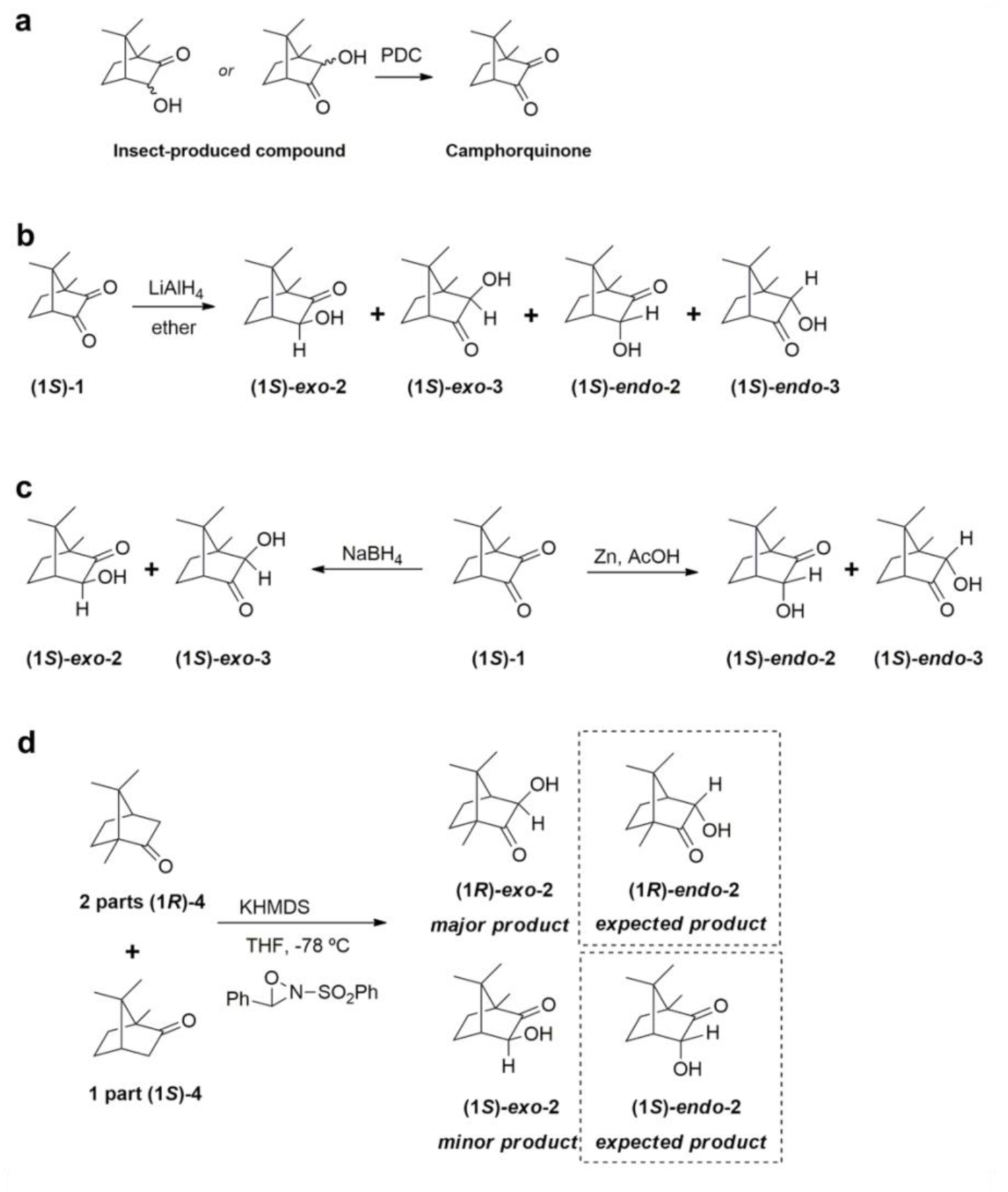
The microderivatization and synthesis experiments used to identify the pheromone compound. (**a**) Oxidation of insect-produced compound with pyridinium dichromate (PDC) yielded camphorquinone **1** as determined by comparison with the mass spectrum and GC retention time of an authentic standard run under identical conditions. (**b**) Nonselective partial reduction of (1*S*)-camphorquinone with lithium aluminum hydride (LiAlH_4_) yielded a mix of four isomers, one of which elicited a response from a male *Photinus corrusca* antenna (Figure 3). (**c**) Reduction of (1*S*)-camphorquinone with sodium borohydride (NaBH_4_), or zinc and acetic acid produced the candidate *exo*- and *endo*-isomers for GC comparison with the insect-produced compound. (**d**) Davis oxidation of a 2:1 mixture of (1*R*)- and (1*S*)-camphor

In a “quick and dirty” proof of the unknown’s structure, partial reduction of (1*S*)-camphorquinone **1** with LiAlH_4_ yielded a mixture of 4 isomers (Figure 2b), one of which matched the retention time of the unknown on achiral and chiral GC stationary phases (DB-5, DB-17, and Cyclodex B), and the mass spectrum of the unknown. Additionally, that isomer elicited a response from the antenna of a male firefly when analyzed by GC-EAD (Figure 3). In total, the data suggested that the bioactive unknown was an isomer of α-hydroxycamphor, likely with the same stereochemistry at the two stereocenters that it shares with (1*S*)-camphorquinone. The remaining uncertainties were the relative positions of the ketone and hydroxyl groups, and whether the hydroxyl group was on the same face as the geminal dimethyl bridge (*exo*), or the opposite face (*endo*), i.e., the four possible isomers shown in Figure 2b. The exact structure, including the confirmation of the absolute stereochemistry, was determined as follows.

**Fig. 3.**
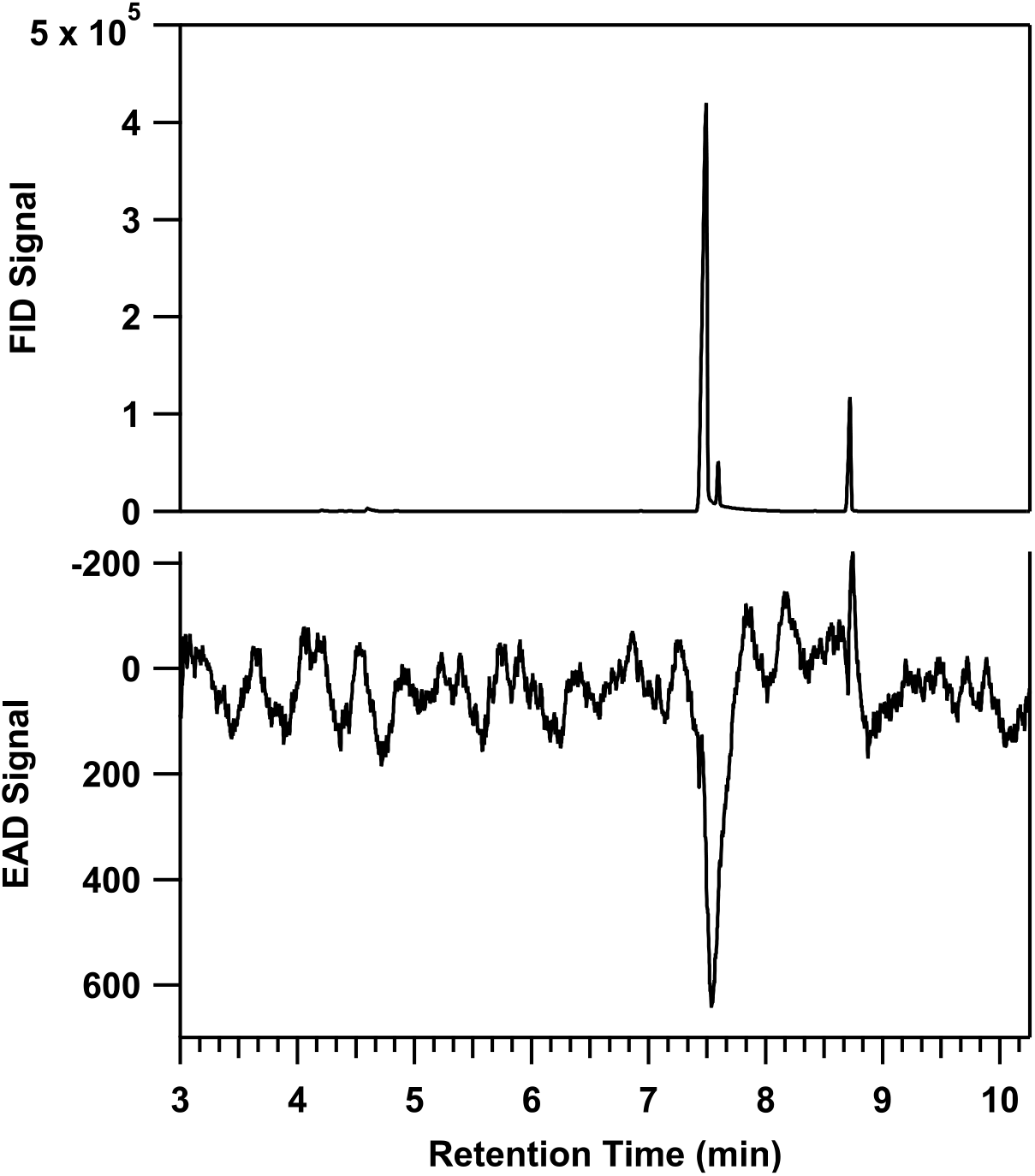
The response of an antenna of a *P. corrusca* male to an isomer from the LiAlH_4_ partial reduction of (1*S*)-camphorquinone

We were fortunate to find established methods for the syntheses of the candidate isomers with sufficient spectroscopic information to identify each one by NMR spectroscopy. To determine the placement of the carbonyl and stereochemistry of the hydroxyl group, mixtures of isomers were synthesized and the individual components were isolated and spectroscopically characterized. To confirm the absolute configuration, commercially available sources of enantiomers of camphor and camphorquinone were used as starting materials to generate products of known absolute configuration. The synthetic standards and the insect-produced compound were then analyzed by GC with chiral and achiral stationary phases (Figure 4) because there was not enough of the unknown in the extracts to isolate and obtain NMR spectra. In these GC analyses, a relatively low injector temperature (120 ºC) was used because of the known propensity for α-hydroxyketones to thermally isomerize (Paquette and Hofferberth 2004). Mixtures of isomers were indeed observed to co-elute when high (240 ºC) injector temperatures were used.

**Fig. 4.**
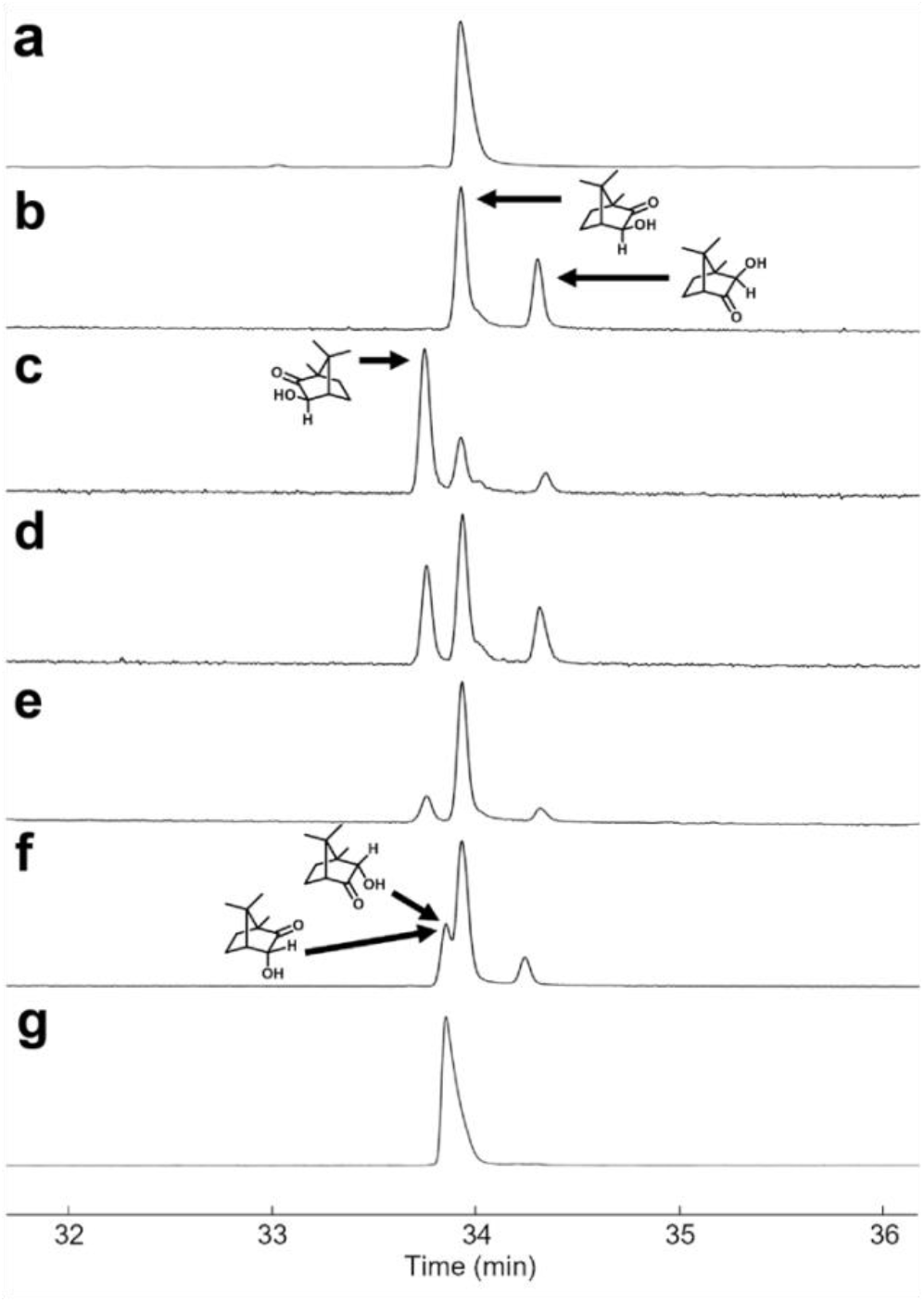
Comparison of synthetic hydroxycamphor isomers to the insect-produced compound. The chiral GC chromatograms illustrate the sequence of injections and co-injections used to identify the gross structure and absolute stereochemistry of the insect-produced compound, and the correct product from the Davis oxidation. (**a**) The single peak shows the insect-produced compound. (**b**) The peaks show the two regioisomeric *exo*-hydroxycamphor isomers (1*S*)-*exo*-**2** and (1*S*)-*exo*-**3** obtained by NaBH_4_ reduction of (1*S*)-camphorquinone, one of which matched the unknown. (**c**) The two *exo*-enantiomers from Davis oxidation of the 2:1 mixture of (*R*)- and (*S*)-camphor (i.e., (1*R*)- and (1*S*)-*exo*-**2**) are well separated, with the minor enantiomer matching the unknown and confirming that the shared stereocenters in the unknown are the same as those in (1*S*)-camphor. (**d**) Injection of the mixture of products shown in panels b and c. (**e**) Injection of the mixture of products shown in panel d and spiked with the insect extract to show the exact match with isomer (1*S*)-*exo*-**2**. (**f**) Co-injection of *exo*-isomers with the co-eluting *endo*-isomers. (**g**) The purified mixture of *endo* isomers (1*S*)-*endo*-**2** and **3** are confirmed to coelute as a single peak, with a retention time different from the insect-produced compound, verifying that the unknown could not be one of the *endo* isomers

To identify which isomer corresponded with which peak in the chiral stationary phase GC traces, product ratios from each synthesis (in accord with literature, see respective synthesis sections) and NMR of the standards (chemical shift, peak integration) were matched with the results from analyses of the same samples on a chiral stationary phase Cyclodex B column (retention time, peak integration). Having unambiguously identified each of the synthetic standards, the insect-produced compound was then analyzed under the same conditions, to match it with one of the isomers.

The first candidates synthesized were the *exo*-hydroxycamphor isomers, by reduction of (1*S*)-camphorquinone ((1*S*)-**1**) with NaBH_4_ (Figure 2c), producing a 65:35 mixture of (1*S*)-*exo*-**2** and (1*S*)-*exo*-**3**. This composition is in accord with the literature (Xu et al. 2006) and allowed for an easy identification of these two isomers in the chiral stationary phase GC chromatogram (Figure 4).

In pursuit of the *endo*-hydroxycamphor isomer (1*S*)-*endo*-**2**, a 2:1 mixture of (*R*)- and (1*S*)-camphor **4** was oxidized with the Davis reagent (2-(phenylsulfonyl)-3-phenyloxaziridine (Figure 2d)(Davis et al. 1984). Two key pieces of information were obtained from this synthesis: (1) the major product formed from the reaction was actually the *exo*-isomer (1*R*)-*exo***-2** (NMR assignment and reaction outcome supported by Piątek and Chapuis (2021) and Neisius and Plietker (2008)) rather than the (1*R*)-*endo*-isomer reported by Davis et al. (1984), and (2) the minor enantiomer (1*S*)-*exo*-2 from (1*S*)-camphor **4** co-eluted with the insect produced compound on the chiral Cyclodex B GC column. Taken together, this indicated that the insect-produced compound was likely to be (1*S*)-*exo*-**2**.

With the two *endo*-hydroxycamphor candidates remaining to be synthesized, a Zn dust/AcOH reduction of (1*S*)-camphorquinone was performed (Figure 2c). This straightforward reaction produced a 56:44 mixture of isomers (1S)-*endo*-**2** and (1*S*)-*endo*-**3**. Even though the identification of each regioisomer on the chiral GC chromatogram could have been difficult because the product mixture is composed of ∼equal proportions, the two isomers coincidentally co-eluted *and* eluted at a different time relative to the insect-produced compound, confirming that neither *endo* isomer could be the insect-produced compound. This allowed us to unambiguously identify the unknown as (1*S*)-*exo*-**2**, formally, (1*S*)-*exo*-3-hydroxycamphor (IUPAC systematic name: (1*S*,3*S*,4*R*)-3-hydroxy-1,7,7-trimethylbicyclo[2.2.1]heptan-2-one).

### Larger Scale Synthesis of Candidate Firefly Pheromone for Field Trials

A larger scale, stereoselective synthesis of compound (1*S*)-*exo*-**2** was carried out by reduction of (1*S*)-camphorquinone **1** with L-Selectride. Unlike syntheses reported in the literature that claimed a completely stereoselective transformation (Kouklovsky et al. 1990; Xu et al. 2002, 2006), an 88:12 mixture of diastereomers ((1*S*)-*exo*-**2** and (1*S*)-*endo*-**2**, 45% yield, inclusive) was obtained after chromatography. Although literature reports claimed that the isomers are inseparable by column chromatography (Xu et al. 2006; Tan et al. 2011), careful flash chromatography on silica gel (20% EtOAc/Hexanes) provided (1*S*)-*exo*-**2** in up to 95% isomeric purity (across multiple experiments), with the remainder being (1*S*)-*endo*-**2**. Finally, the crystalline material was sublimed in a Kugelrohr distillation apparatus, at no detriment to isomeric purity.

### Pheromone-baited Traps Exclusively Attract Male *P. corrusca*

To confirm the biological activity of the candidate pheromone, synthesized hydroxycamphor was deployed in sticky traps at two field locations in *P. corrusca*’s eastern range in May 2022. The first, in Lewisburg, Pennsylvania, was the same site where the individuals used in pheromone collections and GC-EAD had been caught. The second, in Middlebury, Vermont, was near a location where fireflies had been observed in previous years. At both locations, *P. corrusca* males, but not females, were significantly more attracted to sticky traps baited with pheromone lures than to solvent controls (*one-tailed Mann Whitney U test*, PA: P = 0.004, VT: P = 0.004, Figure 5). Not a single male was captured in solvent controls, despite large numbers of males being observed flying slowly around at knee to head height in the habitat (Lower and Holmes, pers. obs. May 2022). Pheromone attraction was confirmed in laboratory assays, where males filmed in an arena spent significantly more time in contact (antennating, mounting, attempting copulation) with the pheromone lure dosed with hydroxycamphor versus the lure with solvent alone (*one-tailed Mann Whitney U test*, P = 0.004; Figure 5).

**Fig. 5.**
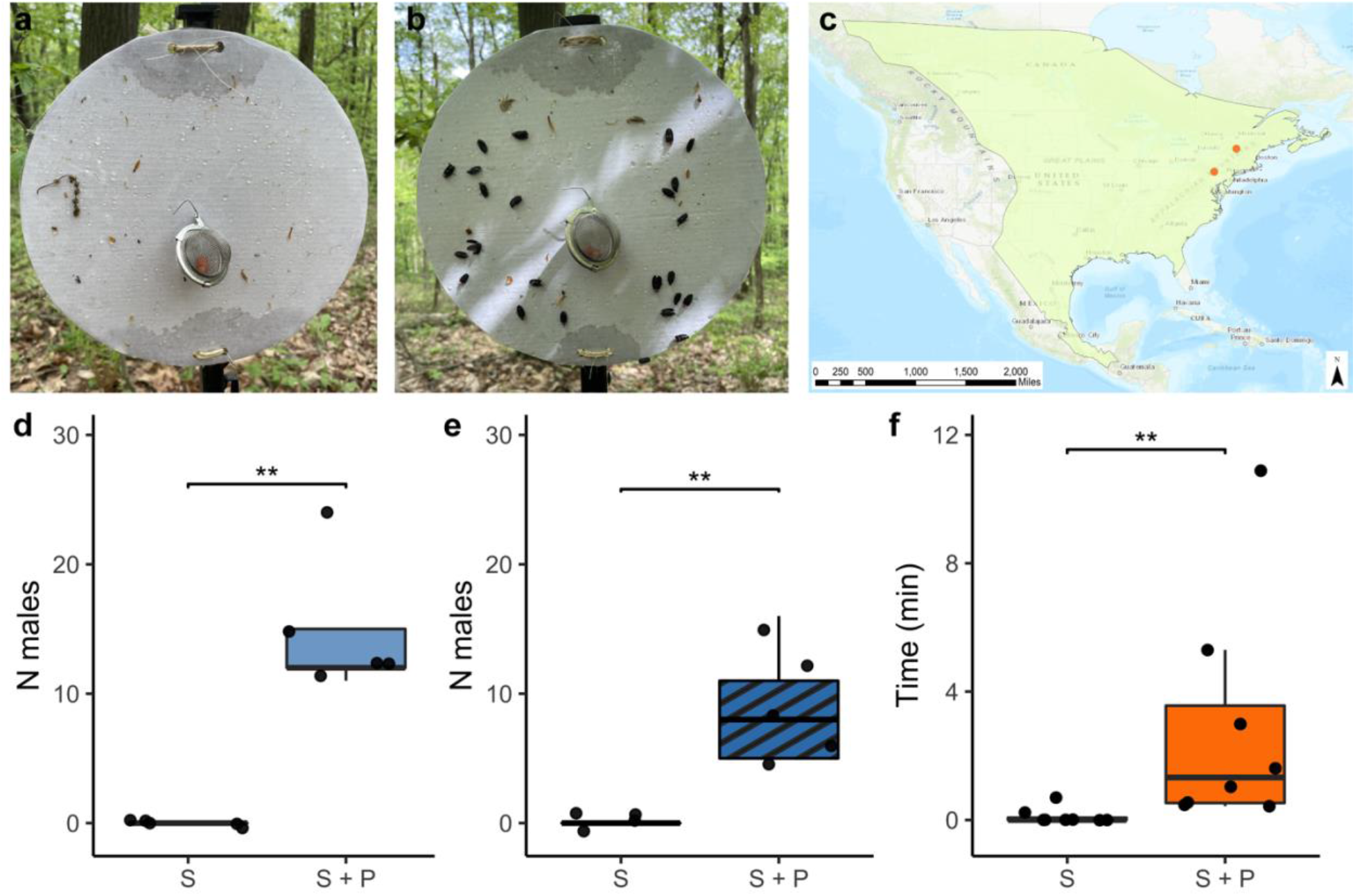
Only male *Photinus corrusca* were attracted to sticky traps baited with synthesized hydroxycamphor. (**a**) Example control (solvent-only) sticky trap with red rubber septum lure visible in the tea strainer after 24 h of deployment mid-May in Lewisburg, PA. While small insect and plant debris by-catch is visible on the trap, indicating the effectiveness of the Tanglefoot, no male fireflies were captured. (**b)** Example baited sticky trap (1 mg dose of pheromone) with trapped male *P. corrusca*. (**c**) North American range of *P. corrusca*. Reproduced, with permission, from Fallon et al. (2021). Orange dots show the two locations where traps were deployed in Lewisburg, Pennsylvania and Middlebury, VT. The range map was created using the World Topographic Map basemap (Esri, 2022) in ArcGIS® software by Esri using data from the IUCN Red List of Threatened Species (IUCN, 2020). ArcGIS® and ArcMap™ are the intellectual property of Esri and are used herein under license. Copyright © Esri. All rights reserved. For more information about Esri® software, please visit www.esri.com. (**d**) Significantly more males were caught in pheromone-baited traps than solvent controls in Lewisburg, Pennsylvania (P = 0.0038, one-tailed *Wilcoxon*). (**e**) Significantly more males were caught in pheromone-baited traps than solvent controls in Middlebury, Vermont (P = 0.0038, one-tailed *Wilcoxon*). (**f**) In laboratory bioassays, males captured in Pennsylvania spent more time in contact with pheromone-treated lures than solvent controls (P = 0.0039, one-tailed *Wilcoxon*)

### Antennal Basiconic Sensilla Respond to Pheromone

We then used a combined approach of SEM imaging and SSR to identify pheromone-sensitive sensilla along the *P. corrusca* antennae. Two morphologically distinct, multiporous basiconic sensilla were identified, with one being slightly shorter than the other (Figure 6a-b, white arrowheads). SSR from the latter sensillum type revealed three olfactory receptor neurons distinguishable by their spike amplitudes. Delivery of puffs of hydroxycamphor elicited dose-dependent responses from the B neuron with the second largest spike amplitude in this sensillum (Figure 6c-d, n=8). The longer multiporous basiconic sensillum (Figure 6a, black arrows) houses two neurons and did not respond to hydroxycamphor (Supplementary Information Figure S3).

**Fig. 6.**
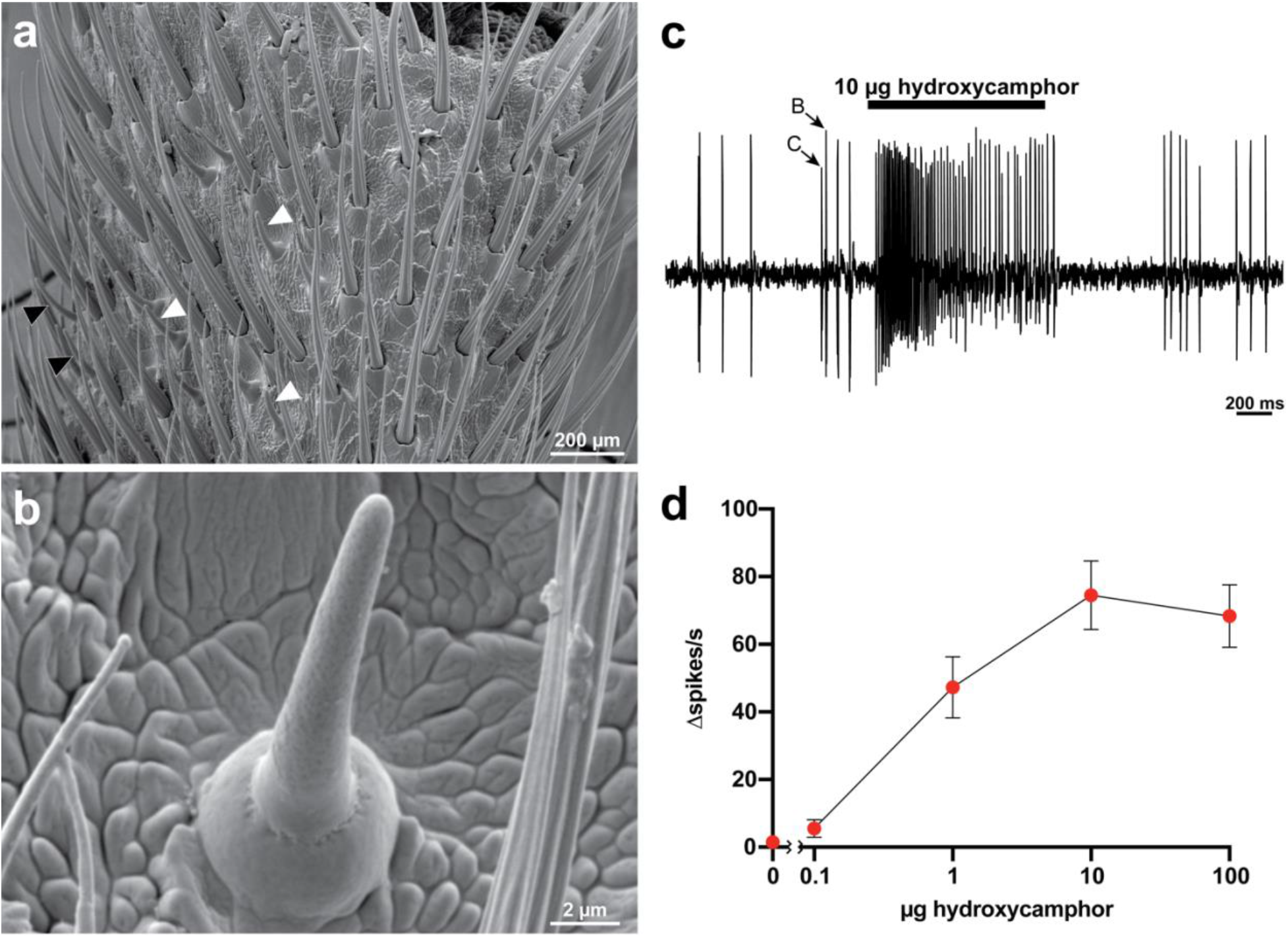
Antennal basiconic sensilla detect *Photinus corrusca* sex pheromone. (**a**) Image of the 8th antennal segment of *P. corrusca*. White arrowheads correspond to basiconic sensilla. Black arrowheads indicate sensilla that were tested, but were nonresponsive. (**b**) Multiporous basiconic sensillum of *P. corrusca* used in SSR experiments of 6C-D. (**c**) Representative SSR recording from a *P. corrusca* antennal basiconic sensillum responding to an air pulse from an odor cartridge loaded with 10 µg of hydroxycamphor. Arrows show spikes from B and C neurons, as identified by differences in spike amplitude and waveform. (**d**) Dose-dependent responses of antennal basiconic sensilla in *P. corrusca* to varying amounts of hydroxycamphor (n=8)

## Discussion

Collectively, our results demonstrate that female-emitted hydroxycamphor (specifically, (1*S*)-*exo*-3-hydroxycamphor) is a sex attractant pheromone of *P. corrusca*. In a previous attempt to identify the sex pheromone of *P. corrusca*, Ming and Lewis (2011) reported that they found significant quantities of *exo,exo*-2,3-camphanediol in whole body extracts of female fireflies. In contrast, we have not been able to unambiguously detect the diol in collections of headspace volatiles across two seasons and two laboratories, whereas the hydroxycamphor pheromone compound was readily detected in most of the headspace samples. It is worthwhile to note that small amounts of camphorquinone were detected in the crude headspace collections, but did not produce a response with GC-EAD (camphorquinone is at RT = 8.72 mins in Figure 3). Our headspace sampling results suggest that the pheromone is not stored in a gland or reservoir, but is likely biosynthesized and released contemporaneously, explaining why it was readily visible in headspace samples collected over a period of days, but was not readily detected in whole body extracts (Ming and Lewis 2011) which would show only a snapshot of what was present at the instant of extraction.

*Photinus corrusca*’s use of a terpenoid as a sex pheromone is not unusual for a beetle (Francke and Schulz 2010). Of the related click beetles that have been studied, many employ terpenoids as female-produced sex pheromones (e.g. Tóth, 2013; Millar et al., 2022). The fact that the pheromone appears to consist of a single component rather than a blend suggests that there is minimal competition for the pheromone channel from congeners, perhaps facilitated by the early-season phenology of *P. corrusca* when few related species are active. The volatile pheromone functions as an effective long-range signal, attracting males over a distance, while visual and close-range chemical signals such as CHCs may take precedence at close range. Whereas in moths, males can be attracted to sex pheromones over hundreds of meters (Cardé and Haynes 2004), in the fireflies *Phosphaenus hemipterus* and *Lucidota atra*, attraction has been reported to occur over shorter distances of ∼20-30 m (Lloyd 1972; De Cock and Matthysen 2005). Our personal field observations during trap deployment suggest that *P. corrusca* males are “bumbling” fliers - generally landing nearby, but not in immediate proximity to the pheromone source. After landing in the vicinity of an emitting female, they actively patrol up and down the tree trunk in search of the source (Lower and Holmes, pers. obs. 2022). Future studies of the range and importance of different signals associated with mating will elucidate their importance in this species.

*Photinus corrusca* is hypothesized to be a species complex rather than a single species (Fender 1970). However, we captured relatively large numbers of males with pheromone traps at two widely separated locations across *P. corrusca*’*s* range, suggesting that if there are cryptic species or subspecies within the lineage, they may still share the same sex attractant pheromone. It is possible that the volatile pheromone could be shared among recently diverged lineages, particularly if they can still hybridize. Future studies using DNA barcoding of samples may offer insights into possible speciation across *P. corrusca* populations.

Recent concern over firefly declines has led to a need for methods to assess population abundance (Fallon et al. 2021). While methods exist for assaying lighted firefly populations using transects (Picchi et al. 2013), photography (Kirton et al. 2012), or video (Sarfati et al. 2020), unlighted species are more cryptic and thus, more difficult to assess. Identification of a pheromone for *P. corrusca* opens up the possibility of using pheromone-baited traps for sensitive and straightforward assessments of presence and population densities of this species. Given that *P. corrusca* represents one of several independent reversions to pheromone-based mate signaling in fireflies (Stanger-Hall et al. 2007, 2018; Sander and Hall 2015; Stanger-Hall and Lloyd 2015; Martin et al. 2017), this also opens the opportunity for future research into the basis of evolutionary transitions in mating signals within this family.

## Acknowledgements

The authors would like to thank the Haussmann family and Bread Loaf View Farm for permission to collect; Sam Pring, Victor Svistunov, and Tobias Ziemke for field assistance and collection; Oliver Keller for morphological insight; and Lawrence M. Hanks and Owais Gilani for statistics consultation.

## STATEMENTS AND DECLARATIONS

### Author Contributions

Sarah Lower spearheaded the project. Sarah Lower, Douglas Collins, Gregory Pask, and Jocelyn Millar proposed and refined the experimental design. Sarah Lower, Gregory Pask, and Douglas Collins acquired funding for the project with Faculty Associate Jocelyn Millar. Sarah Lower collected beetles for pheromone sampling. Sean Halloran performed pheromone collections, and GC-EAD and GC-MS analyses. Yiyu Zheng and Douglas Collins performed aerations and GC-MS analysis for male/female comparisons. Kyle Arriola and Jocelyn Millar identified and synthesized the compound. Hannah Holmes and Sarah Lower conducted field and laboratory assays of the synthesized pheromone in Pennsylvania and performed the statistical analysis, in consultation with Jocelyn Millar. Gregory Pask and Daphne Halley sampled in Vermont and performed the SEM experiments. Gregory Pask conducted the SSR experiments. The first draft of the manuscript was written by Sarah Lower (conceptual overview), Kyle Arriola and Jocelyn Millar (pheromone identification and synthesis), Hannah Holmes (field and laboratory trial methods), and Gregory Pask (SEM and single sensillum recording). All authors assisted with editing the manuscript, and read and approved the final draft.

