## Supplementary Information for "Identification of a Female-produced Sex Attractant Pheromone of the Winter Firefly, *Photinus corrusca*"

Table of Contents

Supplementary Figures 1

Fig. S1 Sticky trap parts and schematic 1

Fig. S2 Terminal segment morphology identifies male versus female *Photinus corrusca* 2

Fig. S3 Responsive and nonresponsive basiconic sensilla on the *Photinus corrusca* antenna 3

References 4

**Supplementary Figures**

**Fig. S1** Sticky trap parts and schematic

Sticky traps consisted of a 10-in white coated-cardboard cake circle with clear packing tape applied to the unlaminated side and edges to increase water repellency, with a metal paperclip folded into a hook inserted through the center. At the PA site, prepared cake rounds (StarMar, Amazon) were tied to the top of 4-foot black fence posts (Parmak, Cole's Hardware) with jute garden twine. At the VT site, prepared cake rounds were fastened to 5-foot light-duty green fence posts (Tractor Supply Company) with plastic zip ties. At both sites, the trap surface was coated with TangleTrap (Tanglefoot, Amazon), leaving a small area in the center clear. After TangleTrap was applied to all traps, a single red rubber septum lure (ACE 9096-32; Vineland, NJ, USA) was inserted into a stainless steel, mesh tea strainer (TIHOOD, Amazon) with the hanging chain removed, and the tea strainer was hung from the paperclip hook in the center of each trap.

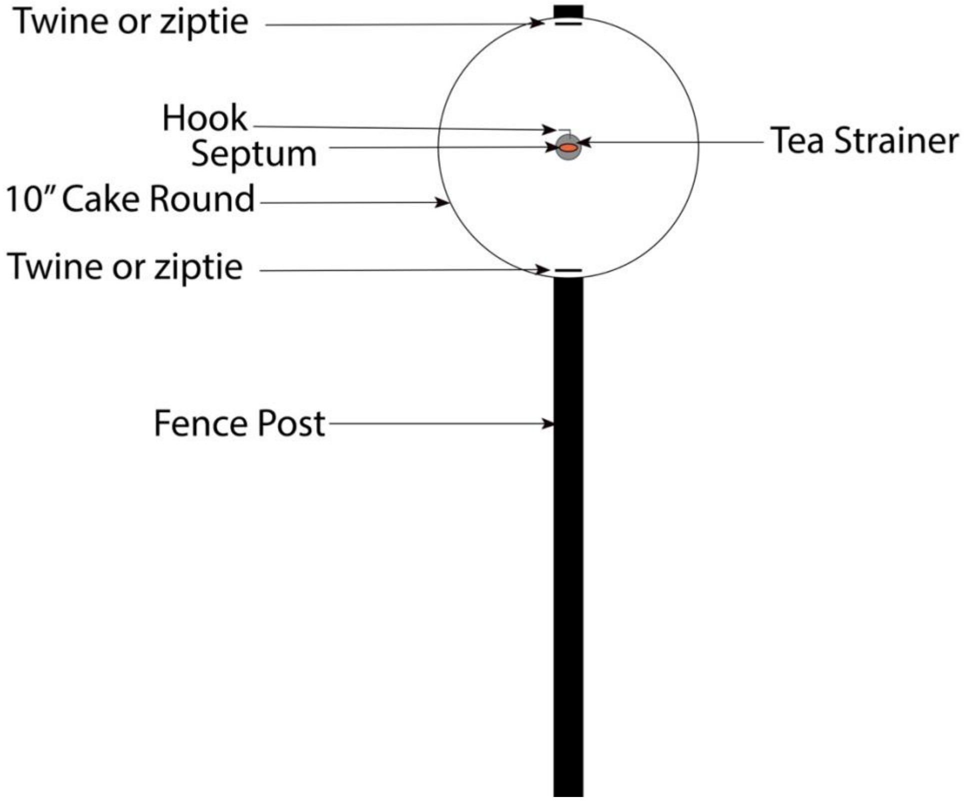

**Fig. S2** Terminal segment morphology identifies male versus female *Photinus corrusca*

Individual *P. corrusca* were identified to sex by examining terminal segment morphology. (A) Females have seven ventrites, with six visible. In females, the sixth ventrite is triangular with a notched tip. (B) Males have eight ventrites, with seven visible. In males, visible ventrite six is sinuous (Lloyd 2002). All individuals caught in sticky traps were confirmed to be male.

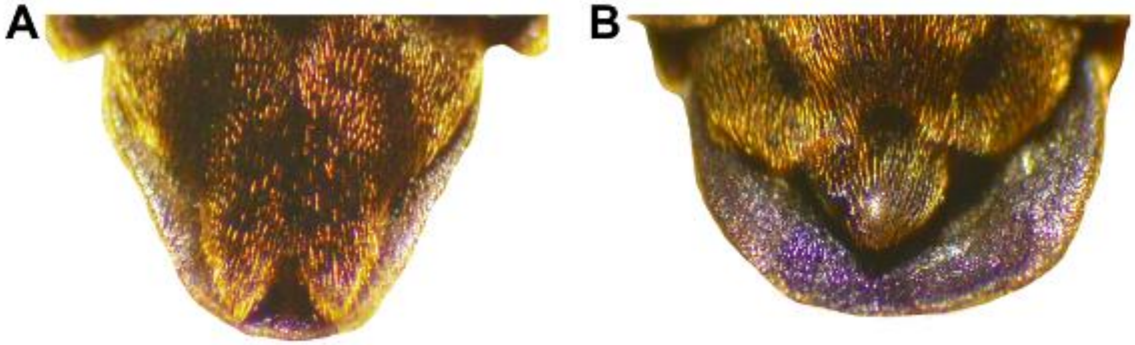

**Fig. S3** Responsive and nonresponsive basiconic sensilla on the *Photinus corrusca* antenna (A,B) Electrophysiological responses of short basiconic sensilla depicted in Figure 6B to either pentane (A) or 100 µg hydroxycamphor in pentane (B) with individual ORN amplitudes differences denoted. (C,D) Electrophysiological responses of short basiconic sensilla depicted in main text Figure 6B to either pentane (C) or 100 µg hydroxycamphor in pentane (D) with individual ORN amplitude differences denoted.

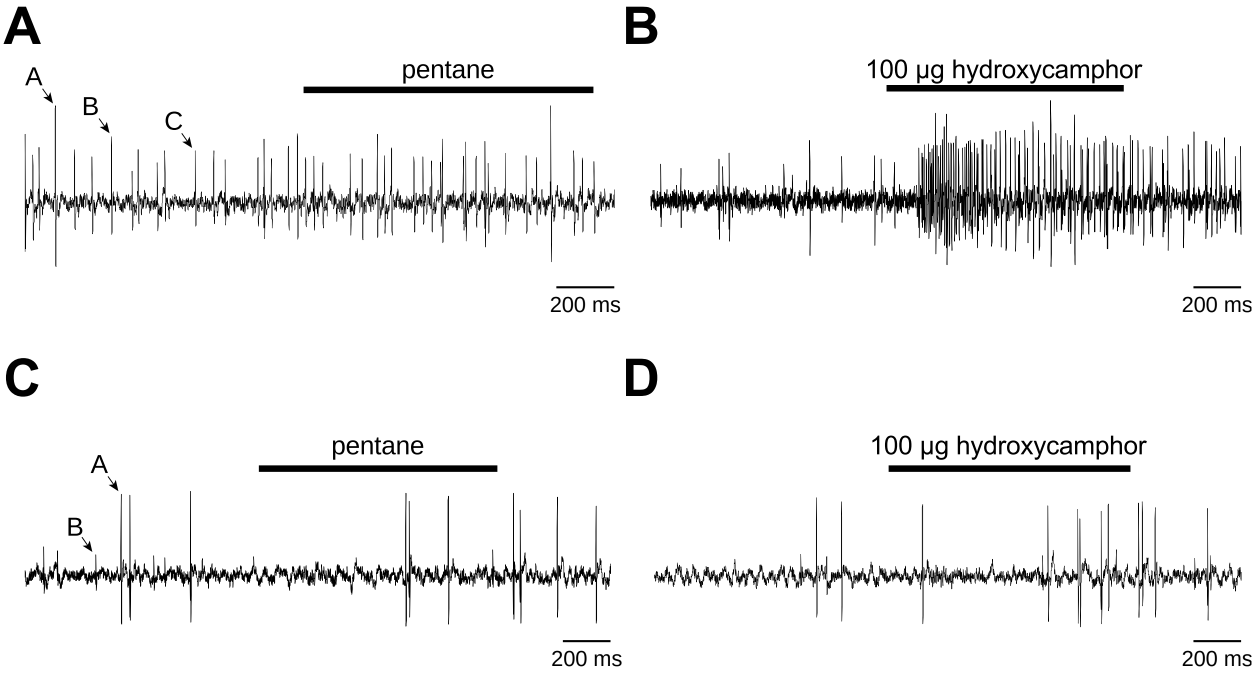
